# A DEVICE THAT COMBINES LIGHT ATTENUATION AND SCATTERING FOR THE MEASUREMENTS OF OPTICALLY DENSE ALGAL CULTURES IN CELL CULTURE FLASKS

**DOI:** 10.64898/2026.09.25.754582

**Authors:** Jose Armando Perez-Loera, Mateo Zavala-Galindo, A. Zavafer

## Abstract

Studying algae requires specialized equipment to characterize culture growth and physiological performance and non-invasive monitoring of these cultures typically relies on optical density (OD) measurements. Currently, OD is often measured using standard spectrometers that suffer from detector saturation at high cell densities, which requires time consuming sample dilutions and lacks compatibility with flat-walled cell culture flasks like Roux bottles. Here we describe a 3D-printable device to measure optically dense algal cultures directly within these cultivation vessels. This instrument combines light attenuation and scattering, resulting in a dual-sensor system that utilizes near-infrared light (950 nm) to precisely quantify biomass without interference from changing photosynthetic pigments. Cultures that reach high cell densities can be assumed to be accurately quantified *in situ* using the scattered signal, preserving the integrity of the culture and eliminating the need for subsampling. This method is adaptable to the most common geometries of Roux bottles (T-25, T-75, and T-175) and could be used to monitor small-scale batch cultures without contamination risks. A cyanobacteria algal species (*Limnospira platensis*) and three flask sizes were analysed on light attenuation linearity and scattering detection limits at different cell concentration, demonstrating that the light attenuation module can reliably measure biomass up to an OD of 3.1 (equivalent to >1.5 g L⁻¹) without requiring dilution.

## 1. Introduction

Studying microalgae often requires specialized equipment to characterize culture growth and physiological performance. Although algae share many experimental needs with more widely studied microbes such as bacteria and yeast, the range of commercial tools designed specifically for algal research is comparatively limited. This gap has led many research groups to develop custom instruments to support their work (Barbosa et al. 2020), and several academic groups have shared open-source designs that expand the set of accessible tools. These open devices are generally not intended to replace commercial systems, which are often more rugged and feature-rich, but instead to broaden experimental possibilities and lower barriers to entry for the community. Many of the available open-source instruments fall into two main categories: cultivation platforms (Erbland et al. 2020; Mapstone et al. 2022; Nedbal et al. 2020; Landschaft and Wishkerman 2024) and tools for assessing biochemical or photophysiological performance (Bates et al. 2019; Nguyen et al. 2022).

There remains a clear need for additional types of sensors and measurement systems to support the diverse requirements of algal research. One of the most widely employed methods for monitoring microbial growth in a non-invasive way is the measurement of optical density (OD), which quantifies light attenuation by cells as a result of absorption and scattering (Sandnes et al. 2006). OD measurements are typically conducted using a transmission configuration, where a sample is positioned between the incident light source and the detector (Oshina and Spigulis 2021). The optical density is expressed as the logarithmic inverse of the ratio between the measured irradiance with a blank and with a sample (Sandnes et al. 2006).

A common use of OD is in growth screening studies in batch cultures (Benner et al. 2022). Typical growth vessels for such studies consist of Erlenmeyer flasks and Roux bottles, and due to the flat walls and short path lengths, the latter vessels can work as low-cost flat panel photobioreactors (Bates et al. 2020). Roux bottles are growth vessels consisting of two transparent spaced flat, rectangular, parallel faces connected to a short neck (Eyre 1913). Their design permits them to keep them standing up or laid down, which makes easier the inspection of the culture on an inverted microscope (Moreau and Furesz 1967; Sánchez-Robles et al. 2015). Despite Roux bottles have high transparency (T > 95%) and their resemblance to optical cuvettes, there are no tools designed to measure optical density in these vessels, either commercially or open source. In contrast, despite the technical complications to measure OD, Erlenmeyer flasks have few open-source tools dedicated to optical measurements in this type of vessels (Mao et al. 2017; Serôdio et al. 2024).

A standard practice is to subsample an aliquot of a liquid culture and measure its OD in cuvettes typically with a 10 mm pathlength (Serôdio et al. 2024). When using common spectrometers and standard cuvettes, good operational practices discourage values exceeding optical density values > 1, as some detectors and digitizers exhibit nonlinear responses at high cell densities, a phenomenon referred to as detector saturation (Yao et al. 2018). However, reaching cell densities that cause saturation is common during most growth experiments, as cell densities often surpass the linear detection range (Yao et al. 2018). Furthermore, at higher concentrations or longer pathlengths, multiple scattering events can dominate, optical density in biological samples breaks the linearity of the classic Beer–Lambert law and introduces nonlinear attenuation (Kocsis et al. 2006). Consequently, when aliquots are measured in standard cuvettes, samples must be appropriately diluted, and the resulting optical density values corrected by the corresponding dilution factor (Myers et al. 2013; Serôdio et al. 2024). This practice can become time consuming when measuring multiple samples at a time or introduce contamination.

Unlike bacterial models such as *Escherichia coli* and *Saccharomyces cerevisiae*, algal cells have a large absorption cross section in the visible spectrum, therefore near infrared (NIR) light is often used to monitor OD (Griffiths et al. 2011). Photosynthetic pigments (e.g., chlorophyll, carotenoids or phycobilins) primarily absorb visible light (Porras Reyes et al. 2024), and these pigments are differentially expressed during the time course of growth. Thus, the determination of OD using light within the visible range can under, or overestimate, biomass (Barbosa et al. 2020). This variation in pigment absorption can yield unstable optical density measurement and to be poor proxy for total biomass (Griffiths et al. 2011). Illumination of 750 nm (OD_750_) has been adopted as an indicator of algal biomass (Clayton 1965; Allen 1968), which contrast to the use of 600 nm which is the typical wavelength for microbial cells like *E. coli* (Yao et al. 2018). Also, OD_750_ was adopted due to signal-to-noise considerations in older spectrometers (Potter and Eisenman 1962; Teich 1970), as they often had higher noise at wavelengths > 780 nm. When OD₇₅₀ nears saturation, measuring optical density at wavelengths greater than 750 nm is recommended, as reduced attenuation at longer wavelengths allows for more accurate assessment of concentration. Hence, optical density determination in the 900 to 1000 nm range can avoiding the need of diluting.

An alternative technique to measure optically dense cultures is the integration of backscattered photon detection when attenuation-based suffer from saturation. This method has been implemented in industrial settings such as pilot plants and fermenters where microbial biomass reaches values of >2 g/ L (Mao et al. 2017). Sensors that integrate both optical density and backscattered signals are highly versatile: attenuation-based OD measurements enable detection of low cell densities, while backscattered light extends the measurable range beyond the saturation limit. But such devices specifically designed for Roux bottles are not commercially available, which would be ideal for lab settings and cultures in small scale.

For all the reasons above, in this work we present a device designed to measure optical density in Roux bottles that combines attenuation and photon scattering for the assessment of grown in microalgal cultures. For the sake of assigning a code to this project, we have named it MODAC (Meter for Optically Dense Algal Cultures). The instrument can be easily manufacture with conventional 3D printers and use of-the-shelf electronics widely available. Despite its small footprint and low cost, this instrument can be used to measure the three most common volumes of commercially available Roux bottles (T-25, T-75 and T-175), making them a useful instrument for phycology labs.

## 2. Materials and Methods

### 2.1 Algal cultures and growth conditions

*Limnospera platensis* (UTEX LB 1926) was grown in Zarrouk media (Phydrotec, the Netherlands) in non-axenic conditions using a bubble-column bioreactor in a greenhouse during the months of November 2025 to May 2026 at the St. Catharines campus of Brock University, ON, Canada (43.1190° N, 79.2490° W). A liter of fresh culture was harvested and diluted to test the MODAC instrument.

Biomass density determinations were done by filtrating 100 mL of culture using a Whatman No. 8 paper filter, which was washed twice with 200 mL of distilled water. Filtration was followed by drying the paper in an oven at 60 °C for two days. To determine biomass density, measurements of the weight before and after the paper filter were used to estimate the total amount of biomass and divided against the isolated volume. The biomass density of the sample used was 1.78 g/L.

### 2.2 Design of the Optical System

The device is constructed around a U-shaped optomechanical scaffold made using 3D printing (Figure 1a), and where light source, detector and samples (Roux bottles) are placed (Figure 1a *a*). The optical sensor is positioned in a cavity indented (Figure 1a *b*) to provide enough space between the bottle and the detecting element. OD measurements use a transmission configuration where a LED (LED_OD_) faces in antiparallel fashion the optical sensor (Figure 1a *c*). The TSL2591 detector board (description of the detector side below) must be placed inside of the cavity, with the jumper cables protruding using the bottom slot (Figure 1a *d*).

**Figure 1:**
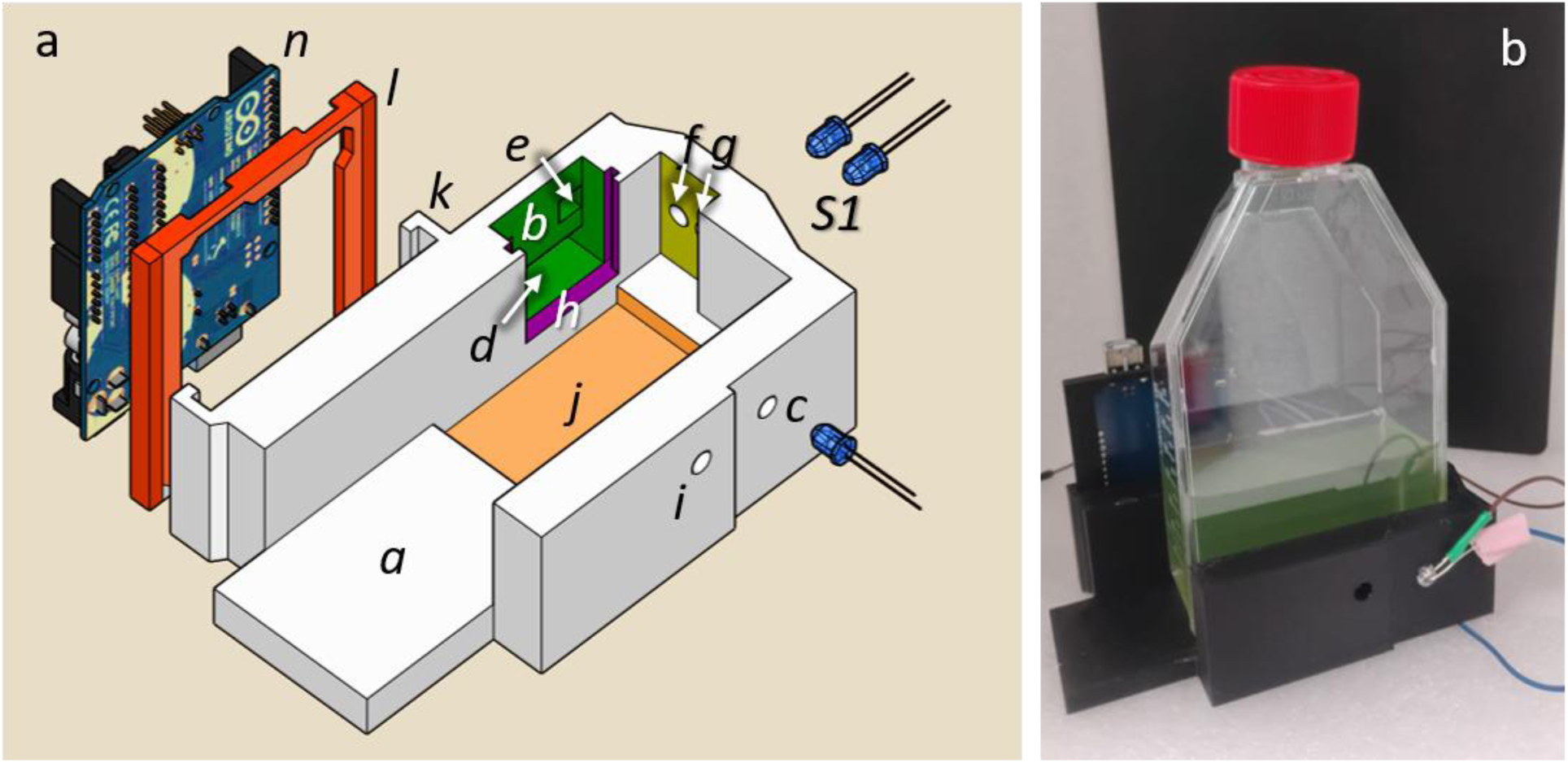
Schematic view of MODAC device and its structure. (A) 3D model used for STL file indicating how to assemble the device. In white, the general scaffold. (a) Sample holder. (b) optical sensor (c) LED for OD measurements. (d) TSL2591 light detector module (description of the detector side below) must be placed inside of the cavity, with the jumper cables protruding using the bottom slot. (f,g). Cavities to install S1 illuminator is resided to prevent the LED lights. (h) Slider at its centre an aperture with the shape of a truncated circle. (i) slot for holding bolt (3 mm diameter, 10 mm length recommended) to hold the bottle in place. (k l) holder for an Arduino UNO board is included. (l) Arduino UNO slider. (n) Arduino UNO (Figure 1A n) and power jacks face upwards. (B) Real MODAC device.

To measure photon scattering, an additional LED is used as illuminator and placed perpendicularly to the sample having effective angles of 114° (S1) (Figure f,g). Cavities to install S1 illuminator is resided to prevent the LED lights from touching the walls of the Roux bottles. The indented optical sensor is protected by a slider at its centre an aperture with the shape of a truncated circle (Figure 1a *h*). This aperture aims to decrease the contributions of off-axis photons during optical density measurements and normalizing the light received by the detector when illuminated by LED S1. All LEDs cavities are width enough to hold standard 5 mm LED in place by friction.

The central space between the detector and the optical density emitter has a width of 41.46 mm and can hold snugly Roux bottles of up to 400 mL (T-175) without scratching the bottle walls. Roux bottles of 60 and 240 mL (T-25 and T-75) can also be placed in the central space, but if additional support is needed on the detector side, a 5.12 mm hole is included to install screws (3 mm diameter, 10 mm length recommended) that hold the bottle in place (Figure 1*a* i).

Adjacent to the optical sensor, a holder for an Arduino UNO board is included (Figure 1a *k*). Printed separately, an Arduino UNO slider (Figure 1a *l* in dark orange) is included in the design and can fit into the side holder where the Arduino USB (Figure 1a *n*) and power jacks face upwards. The enabled device, MODAC, is presented in Figure 1 a *l*, It is advisable to fix the device on to a board or metal sheet to act as a base and counterweight to avoid movement during measurements. All parts were designed using Sketch-up 2025, (Trimble Inc. SketchUp (Version 2024). [Computer software]. https://www.sketchup.com).

### 2.3 Electronics, Firmware and Operation

Light emission was provided by generic transparent 950 nm LEDs connected to digital pins 12 (OD) and 8 (S1) of an Arduino UNO microcontroller (see connection diagram in Figure 2 a). Detection was performed using a TSL2591 dual-photodiode board (Adafruit Industries, NY, USA), which integrates an operational amplifier and a 16-bit analog-to-digital converter (ADC). The firmware was programmed in C using the Arduino IDE. Upon establishing a connection between the MODAC device and a PC, users interact with the instrument via the IDE’s Serial Monitor. Following the on-screen prompts, the measurement protocol proceeds in two steps: first acquiring a blank reading, followed by a sample measurement. After each sample, the MODAC outputs both optical density (OD) and scattering values.

**Figure 2:**
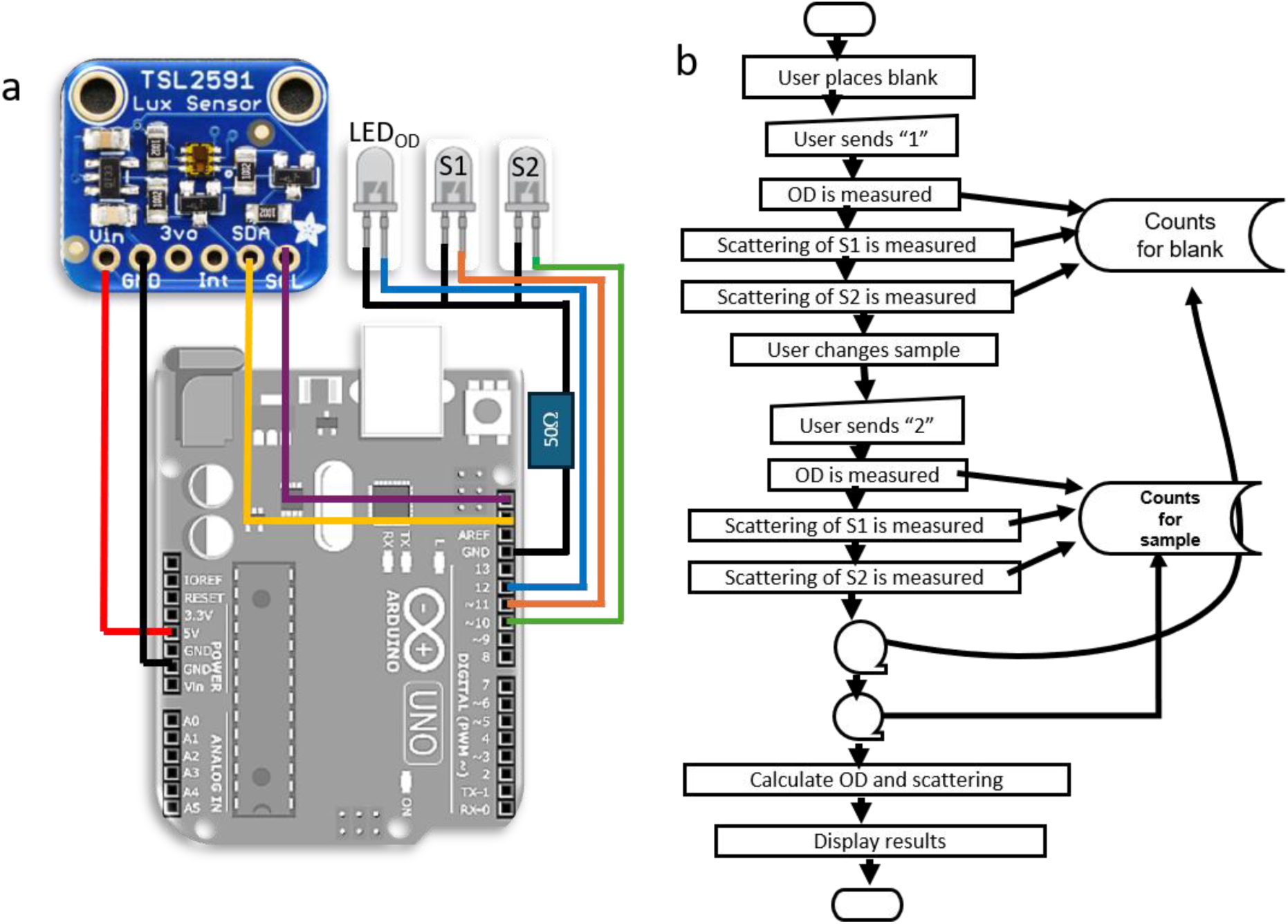
Electronics and firmware diagram. (a) Connection diagram. (b) Flow chart diagram of the operation and data recording.

The measurement sequence embedded in the firmware follows these steps: (1) activate LED_OD_; (2) acquire five readings from the detector and compute their average; (3) store the averaged value in dynamic memory; (4) deactivate LED_OD_; (5) repeat steps 1–4 for LED S1. A flowchart illustrating this sequence is provided in Figure 2b. The TSL2591 module includes two photodiode channels, one filtered for the visible spectrum and one unfiltered. For this application, only the unfiltered channel was used, yielding an effective dynamic range of approximately 40,000 bits. To ensure reliable OD readings at 950 nm, we advise limiting measurements to 3.1 OD. Although a 16-bit system can theoretically reach 4.0 OD, the signal at that level corresponds to fewer than 0 - 10 bits of operational range, making it indistinguishable from noise.

Sensor sensitivity can be adjusted by modifying the integration time, which is selectable across five discrete values (100 to 600 ms). Integration time must be reduced if the transmitted light in the blank exceeds 35,000 counts to avoid saturation. This is particularly relevant in case the blank consists of a slightly light absorbing medium. Firmware and schematics available at: [biophotonicsbrock · GitHub](https://github.com/biophotonicsbrock)

### 2.4 Manufacture and Assembly and of the device

3D modeled parts (optomechanical scaffold, Arduino holder and aperture) were printed using a Creality Ender 3 S1 Pro (Shenzhen Creality 3D Technology Co, Ltd, Shenzhen, China) using generic PLA and slicing the model using UltiMaker Cura ver. 5.6.0 (UltiMaker BV, Utrecht, Netherlands). A custom profile setting (available in our GitHub [biophotonicsbrock · GitHub](https://github.com/biophotonicsbrock)) was used using a 15% infill, 2 walls with 0.8 mm at a 50 mm/s speed and a temperature of 210 °C at the hot end and 60 °C in the hold plate. Tree supports were used in all parts that had a 60° overhangs and brims of 7 mm were used in all models to ensure adequate adhesion. The approximate printing time for the optomechanical scaffold was 3 h, the Arduino holder 20 min and the aperture 6 minutes. TSL2591 was installed using screws and using 3 mm washers by using a Phillips screwdriver and the LEDs soldered and inserted into the scaffold by applying pressure using a flat screwdriver. Having an effective ensemble time after printing of 20 min approximately.

### 2.5 Sample preparation

Fresh spirulina cultures were transferred to 400- (T-175), 240- (T-75) or 60-mL (T-25) polystyrene-made Roux bottles (Sarstedt, Nümbrecht, Germany) using red caps (for adherent cells). Cultures were diluted to create a gradient of cell densities using fresh Zarrouk media.

### 2.6 Spectroscopic Sensor calibration and mathematical model of the instrument

Samples were measured in the MODAC device against a fresh media (blank). Then aliquots were transferred to polystyrene disposable cuvettes (10 mm) and light attenuation was measured using a UV/VIS spectrophotometer (Cary 300, Varian, Melbourne, Australia) at 680(OD_680_), 750 (OD_750_) and 900 nm (OD_900_). (Oshina and Spigulis 2021) If samples displayed optical density values > 1, samples were diluted until optical density values was within range (0 to 1 OD), the final value was estimated by multiplying the sample by their dilution factor (Myers et al. 2013).

Empirically, *OD_l_* was calculated as:

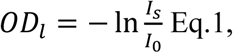

where *I*_0_ was the irradiance measured by the detector for the blank, *I_S_* was the irradiance with the sample. Empirically scattering was calculated by subtracting the counts caused by background scattered photons from the observed scattering by the sample, as done by (Singh et al. 2020; Berne and Pecora 2000).

### 2.6 Data analysis

Data was recovered directly from the Arduino IDE serial monitor and captured into a workbook. All statistical manipulation and plotting were done via Origin Lab Pro 2025. Regression models (linear, single and double exponential) were used to cross-calibrate biomass density with OD and scattering values, their equations are presented in the legend figures. Selection of the model was done empirically by using the most optimum model with the least number of assumptions.

## 3. Results

To confirm that the optical density measured with the MODAC device at wavelengths > 900 nm corresponded with those most commonly used in algal samples we tested the linear relationship with 680 and 750 nm using concentrations of a culture of *Limnospira platensis* (Figure 3a) in a spectrometer. At the three tested wavelengths (680, 750 and 900 nm) the data fit the linear model, and the correlation coefficients are presented in Figure 3b. The main difference between the three tested conditions was the magnitude of the coefficients where 680 nm was double when compared to 900 nm. Using a standard pathlength (10 mm), all wavelengths displayed a linear relationship between concentration OD and therefore had direct equivalence among them.

**Figure 3:**
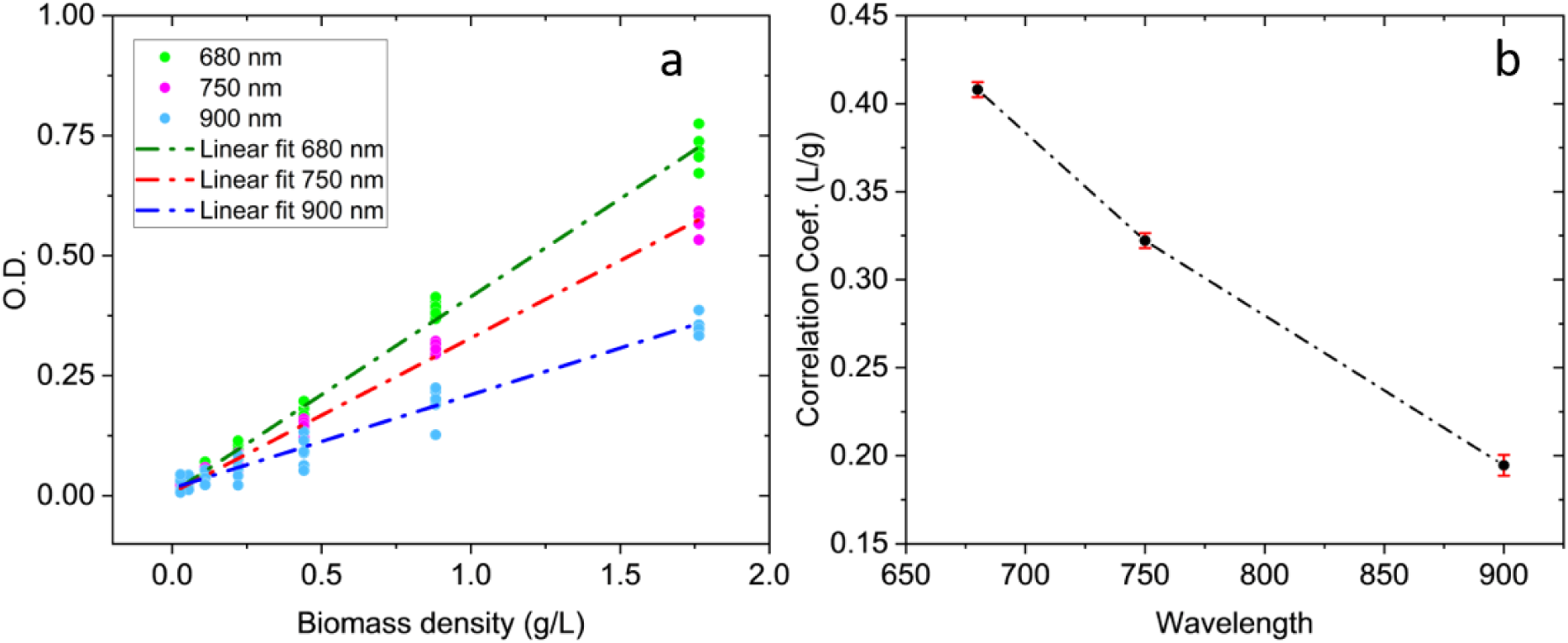
Standardization curves of OD at different wavelengths. (A) linear regression of samples of different concentrations of *Limnospira platensis* using different wavelengths. (B) Correlation values (slopes) of the linear regression +/− standard error. Model used for these regressions was: ***y*** = ***mx*** + ***y*_0_,** where y is the OD, ***m*** is the correlation coefficient, ***x*** is the Biomass density and ***x*_0_** is an offset (this was < 0.015 for all series), r^2^ > 0.98 (for all series).

To assess the linearity and saturation limit for OD measurement obtained with the MODAC device, Roux bottles of different volumes, and thus distinct optical path lengths, were employed at varying biomass concentrations: T75 (Figure 4 ab), T50 (Figure 4 cd), and T25 (Figure 4 ef). In all cases, the relationship between OD and biomass density was best described by a single exponential function fitting well (r² > 0.99). This model provides a robust correction framework for MODAC measurements and acted as calibration curve, enabling precise estimation of biomass density across experimental conditions.

**Figure 4.**
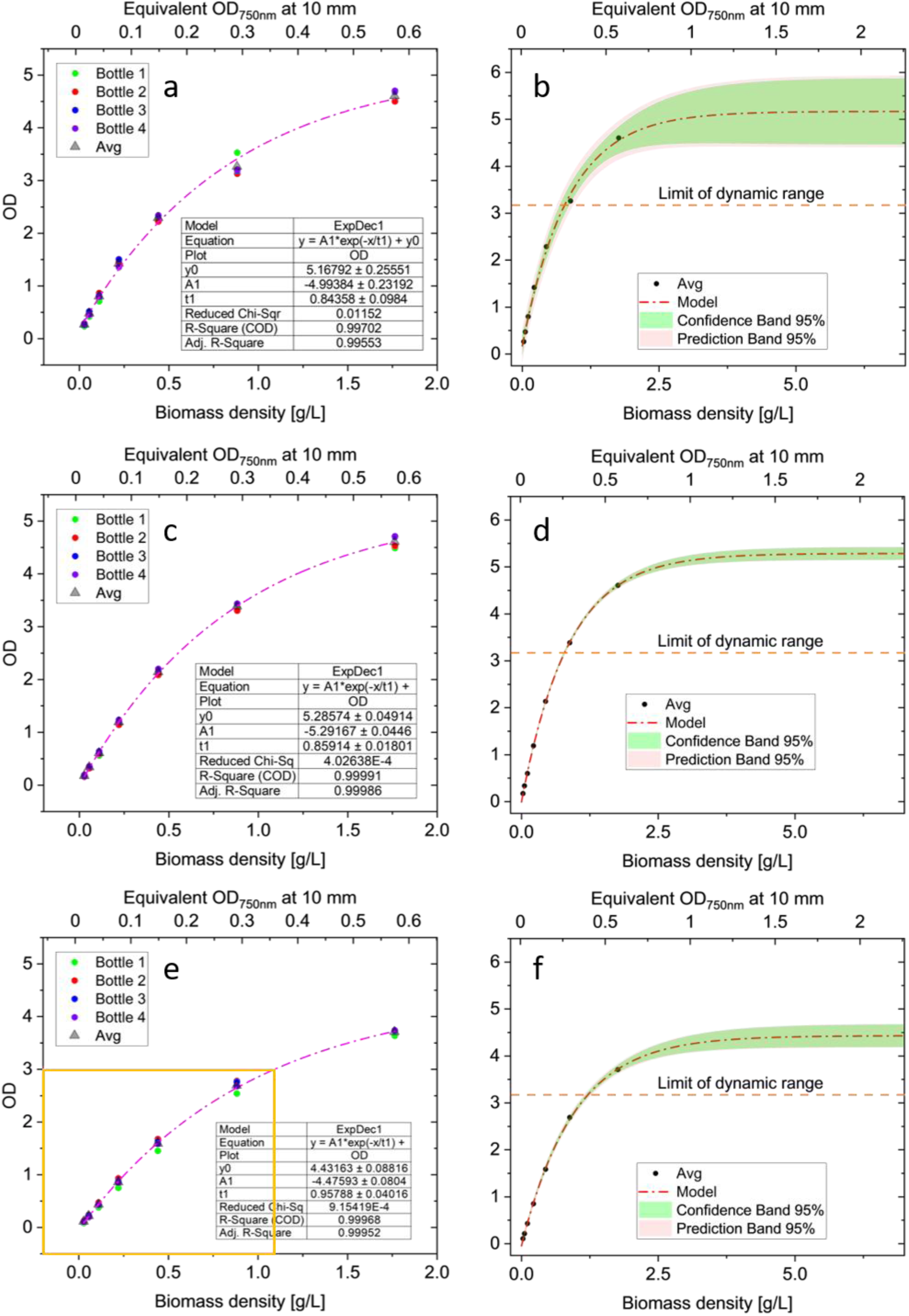
Performance tests of optical density as a function of biomass density of cultures of *Limnospira platensis* using Roux bottles of different volumes. Panels AB display values for T175, panels CD for T75 and panels EF for T25. Panels ACD correspond to the experimental values of four different bottles using different biomass densities, including their respective average and in dotted lines the predictive model is presented. Panels BDF display the model extrapolated to ∞ with their respective confidence and prediction bands of 95%. Orange dotted line represents the limit of dynamic range where the measurements are within the < 5% of actual biomass density. To ease comparison, all plots present the x axis in biomass density (bottom) and equivalent OD at 750 nm using an optical path of 10 mm cuvette in a benchtop commercial spectrometer. Model used for these regressions was: 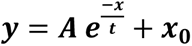, where y is the OD measured in the MODAC device, ***A*** is the amplitude of the calibration curve, ***x*** is the Biomass density and ***x*_0_** is an offset, r^2^ > 0.99 (for all series).

To determine the limit of accuracy for the maximum OD value that the MODAC device can reliably measure, we calculated the number of bits that the TSL2591 module can measure at different OD values and compared them against the exponential model for each bottle including estimation of confidence and prediction of 95% (Figure 4bdf). Since the maximum raw intensity of the IR channel measures is 40,000 bits (OD = 0), a conservative limit of our device are OD values ≤ 3.0, as the dynamic range still will have 40 counts to consider noise of the sample. Under this limit the maximum biomass density for cultures that can reliably be measured is of *L. platensis* are 0.75 g L^−1^ for T-50 and T-175 bottles, while for T-25 can measure up to 1.1 g L^−1^. However, based on the strength of predictive model, for T-50 and T-25 bottles the theoretical limits can be as high as OD = 3.5, as we can predict biomass density with error < 5% (Figure 4df, see prediction bands). Thus, we recommend limiting measurements to a maximum OD of 3.1 when working with highly concentrated cultures.

To complement of the OD measurements, we included measurements of scattered light (Figure 5) and compare them against the same biomass densities evaluated in Figure 4. Scattering in long optical paths, such as the ones of the Roux bottles used here, are caused by multiple scattering events, we opted to fit data to empirical models. For T-25 and T-75 bottles the best fit was a single exponential model, but for T-175 the best empirical model was a double exponential (Figure 5a).

**Figure 5.**
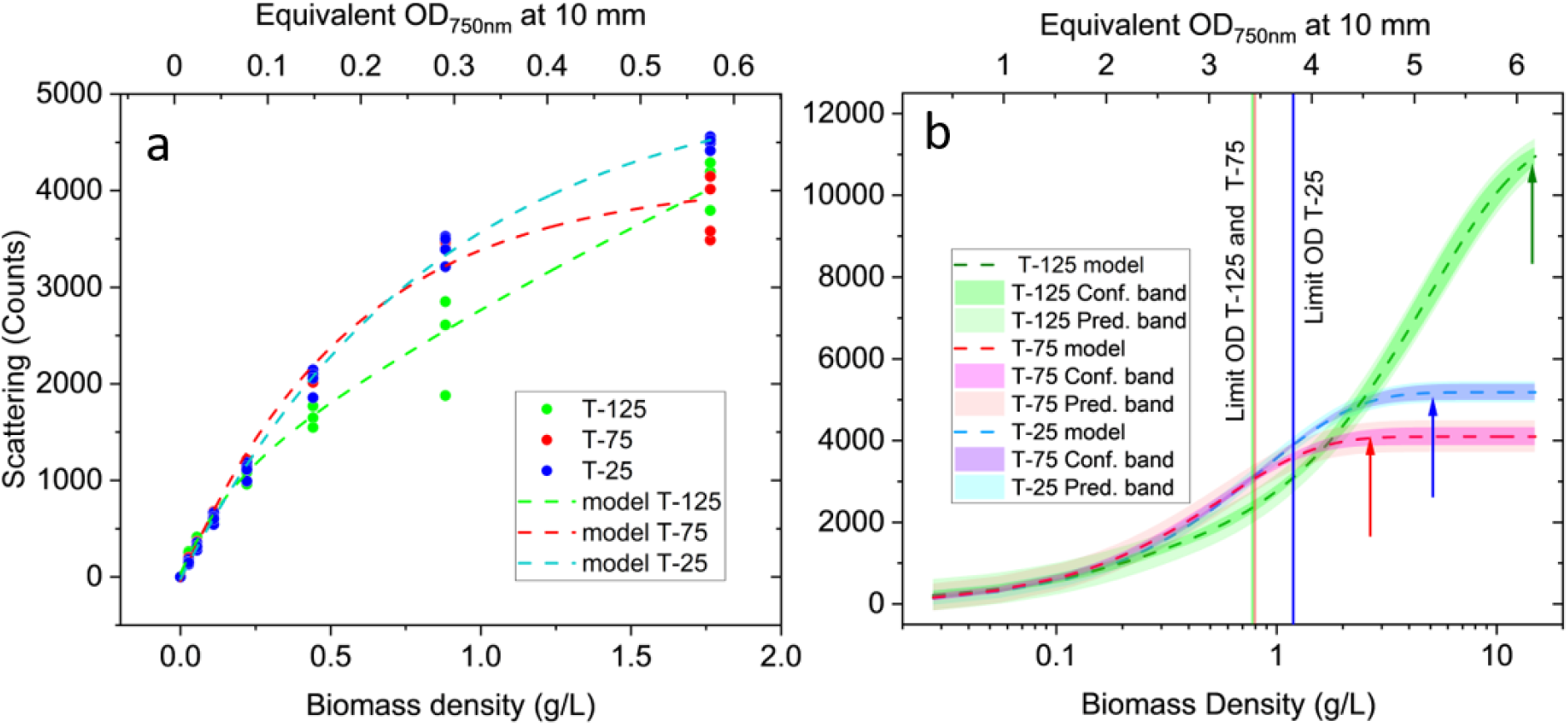
Performance tests of scattering module as a function of biomass density of cultures of *Limnospira platensis* using Roux bottles of different volumes. (A) Experimental values of four different bottles using different biomass densities, including their respective average and in dotted lines the predictive model is presented. (B) Models extrapolated to ∞ with their respective confidence and prediction bands of 95%. Solid vertical lines correspond to the limits of OD measured using the OD module of MODAC and arrows represent the detection limit for each the scattering module. Model used for T-75 and T-25: 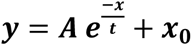, where y is the scattering measured in the MODAC device, ***A*** is the amplitude of the calibration curve, ***x*** is the biomass density and ***x*_0_** is an offset, r^2^ > 0.98 (for the two series). T-175 required a double exponential model 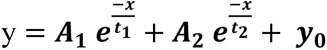, where y is the scattering measured in the MODAC device, ***A*_1_** and ***A*_2_** is the amplitude of the calibration curve, ***x*** is the biomass density and ***y*_0_** is the theoretical saturation point (11,460 bits), r^2^ = 0.98.

The detection range for the scattering module was capable of measuring biomass densities of 1.5 g L^−1^ for the three bottles. Remarkably, only 4,500 bits of a dynamic range of 35 kbits was used. By extrapolating the model to the ∞, we observed that theoretical detection limits for biomass densities of 10, 3 and 5 g L^−1^ are possible for T-125, T-75 and T-25 bottles (Figure 6b). When compared to the biomass density limits of the OD module (Figure 6b), the scattering module extends the detection range of all three types of bottles when compared against OD of the MODAC device (Figure 4bdf).

**Figure 6.**
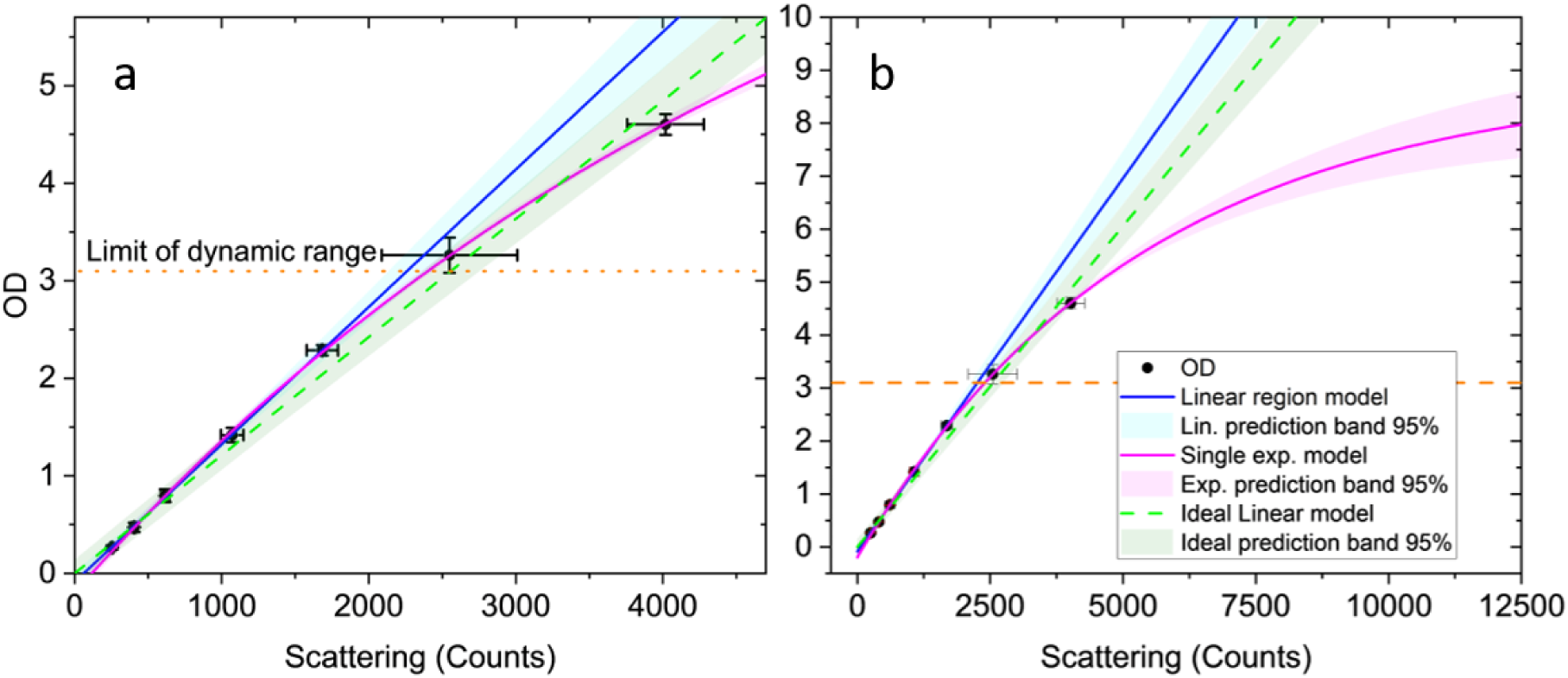
Comparison between OD and scattering values for cultures of *Limnospira platensis* at different concentrations using Roux bottles with a T-175 size. (A) Average of experimental values of four different bottles (± std). The orange line represents the limit of detection for the OD module. Linear model (***m*** = 0.00141 ± 2.68 • 10^−5^ bits, ***x*_0_**= 2.79 • 10^−5^, r^2^ > 0.98) in blue and single exponential model (***A*** = −9.02, ***t*_1_**= 5277 ± 338, ***y*_0_** = **8.81** ± 0.39, r^2^ > 0.99) in pink, shaded areas represent the prediction bands. The green dotted line represents the slope of a linear model in the absence of an offset, presented as a reference point. (B) Models extrapolated to ∞ with their respective confidence and prediction bands of 95%.

Because OD and scattering followed single exponential models for bottles T-25 and T-75, most of light attenuation can be explained due to scattering by biomass in solution. However, in the longest optical path (T-125), scattering required a double exponential model, which means that multiple scattering events are present only when the optical path is long enough for photons to undergo more than one scattering event. To provide some theorical bases to the observations for T-125, we compared OD and light scattering values measured in the MODAC device of cultures. In the range from 0 to 2, OD and scattering of the MODAC device fit satisfactorily to a linear model. However, deviation to linearity is much clearer at greater OD, and the all the datapoints fitting well to a single exponential model. These plateau marks the OD region where the device cannot resolve further increases of biomass, and scattering is a much more effective method to measure growth at high concentrations.

## 4. Discussions

We developed MODAC, a compact, low-cost (USD 20) device for estimating biomass concentration in Roux bottles via optical density. By combining light attenuation and scattering, MODAC can be used to estimate biomass densities even at high concentrations. Assembly takes 20 minutes using only a 3D printer, screwdrivers of two sizes, and a soldering iron, though printing the optical scaffold requires 3 hours using entry-level household SLA 3D printers.

In the example used here, when the two measuring modules are combined, we could measure equivalent values of OD ≍ 3 at 750 nm in 10 mm cuvettes without the need of dilutions (equivalent to biomass densities > 2 g L^−1^). Thus, the main advantage of MODAC is to allow users to measure OD without opening Roux bottles and requiring aliquot subsampling, preventing contamination accidents and saving time in sample handling, much sought needs as noted in (Serôdio et al. 2024).

The integration of LED with wavelengths ≥ 900 nm LEDs into the MODAC device enabled measurements up to 3.1 OD at 950 nm of bottles with longer pathlengths of those of a standard cuvette. Although, we did not explicitly tested 950 nm, previous studies have demonstrated a strong correlation between 950 nm light attenuation and biomass density (Barbosa et al. 2020), giving support to the device presented here. For the model organism used here, *L. platensis*, MODAC can measure equivalent biomass densities > 1.5 g L⁻¹ (dry weight) and 3 to 5 g L⁻¹ (dry weight) when scattering is used. A potential improvement to our device would be the integration of a longer wavelength (e.g., 980 to 1064 nm) which potentially can extend the detection limit to higher cell densities; however, at present this would require a detector module and LEDs/lasers, which are one to two orders more expensive, and were deemed unpractical for the intended design.

A limitation of MODAC is that its scattering response is not linear with biomass concentration, requiring users to generate a species-specific calibration curve before incorporating the device into their experimental workflows. Here, we demonstrate how such a calibration can be constructed and effectively fitted to empirical data. We used *L. platensis* as an example because its filamentous morphology produces a more complex scattering signature than that of simpler unicellular microalgae such as *Chlorella*, *Nannochloropsis*, or *Synechococcus*, all of which are expected to exhibit less complex scattering behavior.

Particularly, the scattering as a function of biomass density can be complex in a square vessel. This is why cylindrical cuvettes (like those used in commercial turbidimeters) are more effective than Roux bottles to be used in backscatter analysis as a curved geometry minimizes edge diffraction and promotes uniform light distribution across the sample (Berne and Pecora 2000).

The incorporation of an illuminator at 117° angles enabled to monitor scattering in Roux bottles with different dimensions, and its position was not decided arbitrarily. The use of different angles in light scattering for the analysis of in microbial cells has been long recognized (Koch and Ehrenfeld 1968). Often, this method is used to monitor particle size, as well of determination of different physical (Berne and Pecora 2000) or biochemical (Narayana Iyengar et al. 2024) properties of cells. Most notably, light scattering is widely used in flow cytometry (Kerker 1983) to monitor different population sizes of cells or organelles (Steen 2004). However, such uses require a collimated light source, such as a coherent source, thus, we advise against using our devices to extract information beyond biomass density using the scattering channel. This is because LEDs are anisotropic (Marcato et al. 2025; Mehta et al. 2010), non-collimated (Chen et al. 2012) and incoherent light sources (Malacara-Hernández 2017; Mehta et al. 2010) with a highly divergent beam (Mehta et al. 2010) incompatible to compare against an actual angle required for particle determination.

There have been other open devices to measure optical density in microbial cultures to name few, but conceptually they are very different to the device presented here, as they have designed for flow through operation, microfluidics, etc. (Jia et al. 2015; Yao et al. 2018; Nguyen and Rittmann 2018). The MODAC adds to collection of open-source tool to study algae grown in conventional vessels like media bottles (Parret et al. 2025), Erlenmeyer flasks (Serôdio et al. 2024) and well plates (Hasson and Wishkerman 2022).

It is fair to indicate that our device draws inspiration of two previous open projects, in particular the *Phenobottle* (Bates et al. 2020) and *Erlenmeter* (Serôdio et al. 2024). The *Phenobottle* is an open-source automated photobioreactor to study the photophysiological characterization of microalgae in 240 mL Roux bottles. Its shape inspired the shape of our optomechanical scaffold. In contrast with our system, the *Phenobottle* measures optical density using two identical 875 nm LEDs, one as a light source and another one as a detector, connected to the microcontroller via a pull-up resistor acting as a gain. Like the *Phenobottle*, our device requires a microcontroller as DAQ, but because the measurements do not need high processing speeds like the microcontroller used in the *Phenobottle*, our device was designed round the Arduino UNO. Furthermore, the Teensy 3.6 boards (microcontroller used in the Phenobottle) are no longer manufactured and their replacement exceeds the cost-benefits when compared when an Arduino microcontroller and the TSL2591 detector board that is equipped with a 16-bit ADC.

While one of our initial prototypes considered a similar optical density module used in the *Phenobottle*, a TSL2591 includes an OPAMP and has from factory a linear response across its dynamic range, making it more practical. This decision was influenced by the *Erlenmeter*, which is another low-cost, open source turbidimeter for Erlenmeyer flasks (Serôdio et al. 2024). Despite challenges to measure optical density in conical flasks (Mao et al. 2017), the Erlenmeter notably maintained linearity up to OD_466_ 1.25 equivalent in 10 mm cuvettes (Serôdio et al. 2024). But unlike Erlenmeyer flasks, Roux bottles have an even optical path and the optical density can be measured in any part of the bottle. The magnitude of OD is directly proportional to the path decreasing the risk of inconsistent and unreliable readings. All good characteristics when experimental needs require to monitor growth at high cell densities.

Another major difference with the *Erlenmeter* is that we opted for the use of a 950 nm detection light instead of any illuminator in the visible range to avoid interference from pigment absorption by chlorophyll *a* and *b* (Griffiths et al. 2011). Although, this is optionally to be equipped in case users requires to do so. However, users should be aware that measuring at 466 nm means the measured optical density is a combination of light scattering by the cells and light absorption by the chlorophyll (Griffiths et al. 2011). As the algae grow, both factors may change, decreasing the likelihood to isolate the signal due to biomass (cell concentration) (Griffiths et al. 2011) from the signal due to physiological changes (pigment content) (Mohsenpour et al. 2012). Changes in the algal culture’s physiological state (e.g., nutrient stress, light-dark cycles) can cause the chlorophyll content to vary significantly, even if the cell number remains relatively constant (Griffiths et al. 2011). Another advantage of the use of 950 nm is not photochemically active in algae which allows the analysis of cells without disrupting dark adaptation which is relevant for chlorophyll fluorometry (Kalaji et al. 2014) or dark respirometry (Xue et al. 1996).

In summary, MODAC is a cost-effective system to monitor algal growth in Roux bottles of different sizes. The OD module allows users to monitor in a conventional way growth but saturates at lower biomass densities when compared to scattering. Scattering in the other hand allows to extend detections. The scattering module proved to have at extended range in bottles with longer path lengths and it is recommended to be used at high biomass densities.

Over the last decade multiple open and semi-open-source optical density sensors have been published showing an active community in the field of microbial bioengineering. Often each design aims to be implemented a specific purpose (pigment detection, biomass accumulation, strain specific needs, etc.) or accompany the design of a new type of bioreactor. A common theme has been the use of flow-through systems for the in-line monitoring of cultures in continuous mode, and the offering for tools for the study of batch cultures in small scale has been significantly shorter, so we hope the low cost and ease of construction of the instrument presented here will help labs expand their capabilities.

## Declarations

### Availability of data and material

Data available at Federated Research Data Repository https://www.frdr-dfdr.ca

### Author’s contributions

#### CRediT statement

Conceptualization: AZ; Methodology: APL, AZ; Software: AZ; Validation: APL, MZ; Formal analysis: AZ; Investigation: APL; Resources: MZ, AZ; Data Curation: APL AZ; Writing - Original Draft: AZ; Writing - Review & Editing: AZ, APL ; Visualization: AZ; Supervision: AZ; Project administration: AZ; Funding acquisition: AZ.

## Acknowledgements

The team would like to thank the technical support and training of Margarita Di Profio, Irene Palumbo, Alison Smart and Dr. Aditi Das.

